# High-speed multifocus phase imaging in thick tissue

**DOI:** 10.1101/2021.06.16.448670

**Authors:** Sheng Xiao, Shuqi Zheng, Jerome Mertz

## Abstract

Phase microscopy is widely used to image unstained biological samples. However, most phase imaging techniques require transmission geometries, making them unsuited for thick sample applications. Moreover, when applied to volumetric imaging, phase imaging generally requires large numbers of measurements, often making it too slow to capture live biological processes with fast 3D index-of-refraction variations. By combining oblique back-illumination microscopy and a z-splitter prism, we perform phase imaging that is both epi-mode and multifocus, enabling high-speed 3D phase imaging in thick, scattering tissues with a single camera. We demonstrate here 3D qualitative phase imaging of blood flow in chick embryos over a field of view of 546 × 546 × 137 μm^3^ at speeds up to 47 Hz.

## 1. Introduction

Phase microscopy is widely used to image intrinsic index-of-refraction variations in biological samples [1]. Since it does not require external staining or labeling, transparent cells or organisms can be observed in their natural states. Over the past decades, many techniques have been developed, both qualitative and quantitative. Techniques based on incoherent sources such as lamps or LEDs include differential interference contrast [2], lateral shearing interferometry [3], illumination/detection pupil modulation [4–7], and numerical phase retrieval from image defocus [8–10]. Techniques based on laser illumination include digital holography [11], quadrature interferometry [12], and spiral phase contrast microscopy [13]. However, a common drawback of all these techniques is that they fail to provide depth discrimination: they provide only 2D phase maps obtained from either the microscope’s focal plane or from axial projections (integrations) through the entire sample. Because many biological samples present index-of-refraction variations that naturally vary in 3D, it is often desirable to image these variations over extended volumes.

Several techniques have been proposed to reveal phase information in 3D. The most popular of these is optical diffraction tomography, where measurements of the scattered light intensity [14–16] or complex field [17–20] under different illumination angles are used to recover the 3D sample phase, under the assumption of single scattering. By incorporating more complex numerical models, this strategy can be extended to multiple scattering in thick samples [21–23]. Alternatively, 3D phase information can also be retrieved from 3D focal stacks, typically obtained by means of physically translating the sample/camera [24–26] or remote focusing [27]. Because these techniques involve many sequential image acquisitions, their applications are generally limited to static or slowly varying samples.

To observe fast dynamic samples in 3D, more recently, single-shot 3D phase imaging has been demonstrated by using custom-fabricated beamsplitters [28–30]. In this strategy, multiple focal planes are simultaneously imaged with one or two cameras without axial scanning. 3D depth-dependent phase information becomes intrinsically apparent in the 3D focal stack with the use of oblique illumination [29, 30], or can be retrieved *post hoc* by numerical deconvolution [28]. Because volumetric information is captured in a single shot at the camera frame rate, fast events such as cilia beating can be observed in real time. However, because these techniques rely on the use of trans-illumination, they are restricted to biological samples that are thin and near-transparent.

In this work, we introduce multifocus oblique backillumination microscopy (multifocus OBM) for high speed 3D phase imaging in thick, scattering samples. Our technique is based on a previously developed strategy [31, 32], where multiple scattering from the sample is used to convert off-axis epi-illumination into oblique trans-illumination, thus enabling optically-sectioned phase-gradient contrast in arbitrarily thick samples [32, 33]. To enable this technique to record volumetric information, we make use of a z-splitter prism [29] that allows up to 9 distinct focal planes to be captured simultaneously with a single camera. By combining instantaneous focal stacks obtained from a rapid sequence of 3 different azimuthal illumination angles, 3D qualitative phase images are then constructed by a method of complex Fourier integration [34]. We demonstrate our technique by imaging the chorioallantoic membrane (CAM) of chick embryos over a volume of 546 × 546 × 137 μm^3^ at a speed of 47 Hz, limited by our camera frame rate.

## 2. Methods

### 2.1. Multifocus oblique back-illumination microscopy

The principle of OBM is detailed in [31]. Briefly, OBM relies on tissue scattering to create oblique trans-illumination in an epi-illumination geometry. When light is delivered into the tissue from the surface, it becomes multiply scattered and a portion of this light is directed back toward the sample surface. If the illumination is delivered off-axis, this results in an overall off-axis tilt of the backscattered illumination that transmits through the focal plane from below, which, in turn, enables the possibility of phase-gradient contrast when imaging through a microscope objective.

In the initial implementation of OBM, two diametrically opposed off-axis fiber light guides were used to separate phase-gradient contrast from absorption contrast at the 2D microscope focal plane [31, 35]. Extended-depth-of-field phase-gradient imaging beyond this focal plane was then demonstrated by using a fast scanning electrically tunable lens [32]. More recently, full 3D quantitative refractive index characterization was achieved by deconvolving 3D OBM stack images using a numerically-modeled phase transfer function [36]. In this last strategy the image stacks were acquired by physically translating the sample, precluding the possibility of real-time imaging.

To enable fast volumetric imaging of 3D phase information, we combine here the principle of OBM with a recently developed multifocus imaging strategy based on a z-splitter prism [29]. Our experimental setup is shown in Fig. 1 (a), which is similar to the multifocus microscope described in Ref. [29] but with illumination delivered through off-axis optical light guides. In brief, illumination light from LEDs (Thorlabs, M617L3) is coupled into 1 mm diameter plastic light guides (Edmund Optics, #02-536), and delivered into the sample at a 2.5 mm offset distance from the optical axis, and at a roughly 45° angle relative to the sample surface. To enable the robust extraction of phase gradients in both transverse directions, we used three azimuthal illumination directions separated by 120° [Fig. 1 (b)]. The optical light guides are affixed under the objective by a custom 3D-printed holder. For detection, the transmitted light is collected by an objective lens and focused by a tube lens *f*_1_ through the z-splitter prism to create multiple discrete focal planes that are then re-imaged by an additional 4 *f* relay system to a common large-area sCMOS camera (Hamamatsu ORCA-Flash 4.0 V3 – full frame rate 100 Hz) for recording. During imaging, multifocus phase-contrast image stacks from the three azimuthal angles were collected sequentially by externally triggering both the camera and LEDs with a multi-function DAQ board (National Instruments PCIe-6321).

**Fig. 1.**
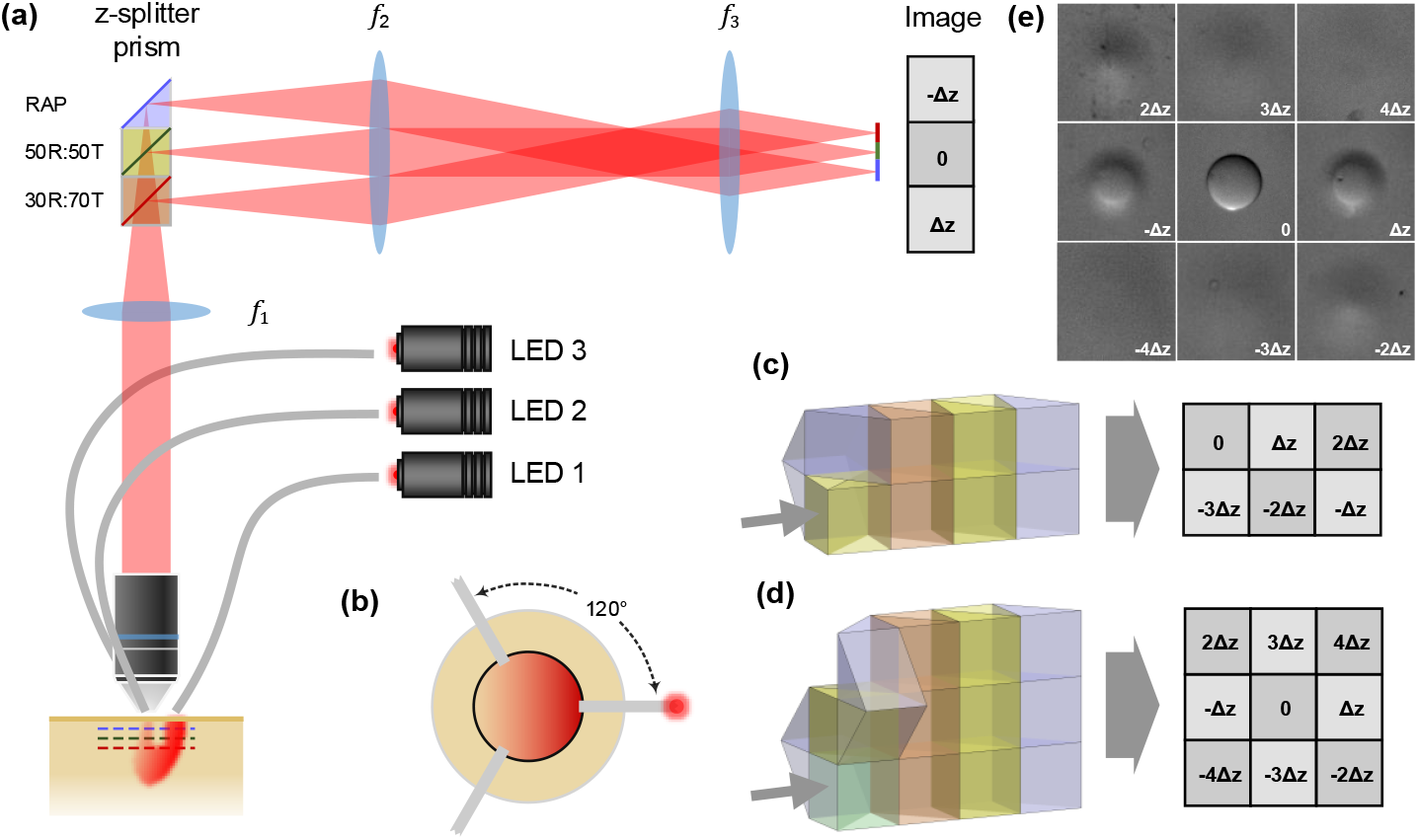
(a) Multifocus OBM setup with a 3-plane z-splitter prism. *f*_1_ = 200 mm, *f*_2_ = 300 mm, *f*_3_ = 100 mm. (b) Illumination with 3 different azimuthal angles separated by 120° through 3 individual optical light guides. Constructions of z-splitter prisms for (c) 6-depth imaging, and (d) 9-depth imaging. Orange cube, 30R:70T BS; yellow cube, 50R:50T BS; green cube, 70R:30T BS; purple cube, RAP. (e) Single-shot 9-depth images of a single 45 μm polystyrene bead with illumination from a single azimuthal angle. Δ*z* = 29.5 μm.

The key to our multifocus imaging system is the z-splitter prism. In our case, this was obtained from Artifex, though alternatively it could be assembled entirely from off-the-shelf beamsplitters (BS’s) and right angle prisms (RAPs) [29]. The functions of these is to split the detection path into multiple paths of increasing optical pathlength, so that each path, when projected onto the camera, is conjugate to a different focal plane in the sample. The size (separation distance) *L* of each BS/RAP determines the axial interplane separation Δ*z*:

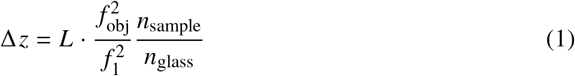

where *f*_obj_ is the focal length of the objective, and *n*_sample_ and *n*_glass_ are the refractive indices of the sample and the BS/RAP glass respectively. Z-splitter prisms can be made to image 3, 6, or 9 distinct focal planes by using different BS/RAP combinations. In this work we used a 9-plane z-splitter with BS/RAP dimension of *L* = 12.5 mm, material N-BK7 glass (*n*_glass_ = 1.516), which we utilized in either a 6- and 9-plane imaging configuration depending on the entrance port into the prism.

As an example, we imaged a single 45 μm polystyrene bead embedded in PDMS from a single oblique backillumination angle. The simultaneously captured 9-depth images are shown in Fig. 1(e), where the interplane separation is Δ*z* = 29.5 μm. Since polystyrene beads exhibit very little absorption, the raw OBM images primarily contain phase contrast. By virtue of the optical sectioning that comes inherently from phase-gradient imaging [31], the bead is most visible only near Δ*z* = 0.

### 2.2. Qualitative phase image reconstruction

For most biological samples, images from a single oblique illumination angle contain contrast from both phase and absorption. One must generally take at least two images of opposing tilt angles to isolate phase-gradient contrast from absorption contrast. Once pure phase-gradient contrast is isolated it is then possible to reconstruct qualitative or quantitative phase images. Here we use a similar procedure to reconstruct multifocus images.

We start with a general formalism for OBM images acquired with *N* different illumination azimuthal angles evenly distributed over 2*π* (in our case *N* = 3). We denote *I_z,n_* as the raw OBM images at depth Δ*z* = *z* from illumination angle 2*nπ/N*, where *n* = 0, 1*, …, N* − 1. For each depth, the absorption contrast is simply the sum of these images

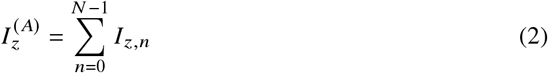

and the phase-gradient contrast along the transverse axes *x, y* can be synthesized from:

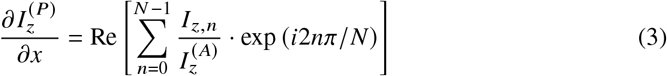

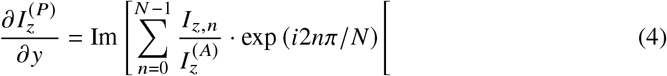

where Re[·] and Im[·] are the real and imaginary parts of the associated argument. A number of methods can be used to retrieve phase images from phase-gradient images. Here we use the method of complex Fourier integration since it is highly robust [34]. In this technique, a complex gradient image is first defined as

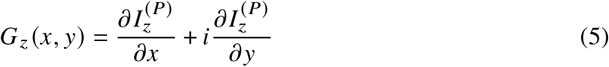

The phase image is then obtained from:

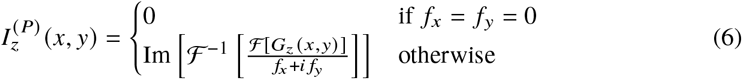

where 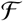 and 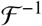 are Fourier and inverse Fourier transforms, and *f_x,y_* are the corresponding spatial frequencies. This procedure is repeated for all individual z planes, allowing us to construct a qualitative 3D phase image stack, as illustrated in Fig. 2.

**Fig. 2.**
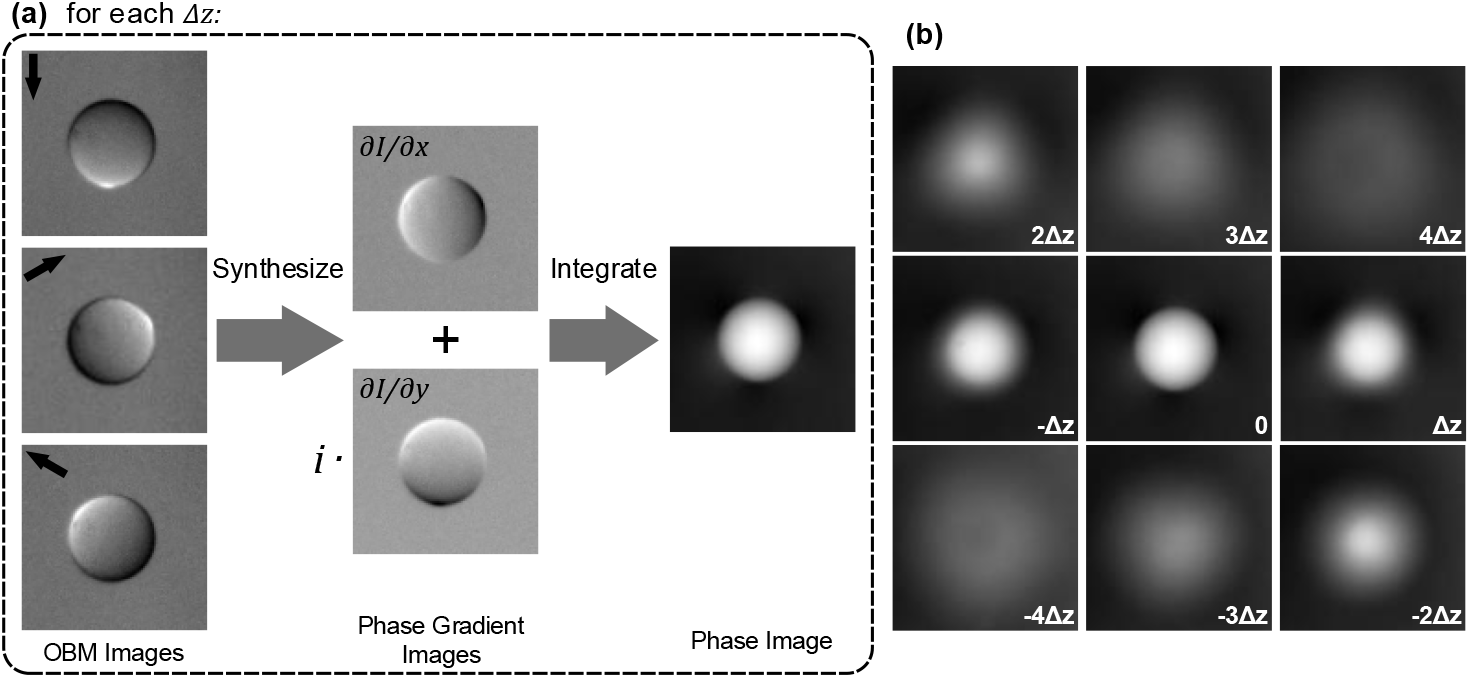
(a) Procedure for reconstructing qualitative images from raw 3-angle OBM images. (b) Reconstructed 9-depth qualitative phase image of a single 45 μm polystyrene bead. Δ*z* = 29.5 μm.

Prior to constructing phase images, the raw images from each depth were cropped and registered according to [29]. To correct for illumination non-uniformity, each raw image was divided by a Gaussian-blurred (radius = 50 pixels) version of itself, effectively flattening them.

As a side note, for phase reconstruction, we adopted a 3-angle illumination strategy enabling phase gradients to be synthesized in both transverse x,y directions. A similar strategy applied to oblique detection using a pyramid wavefront sensor has been studied as well [37, 38]. Compared to other OBM implementations that use 4 opposing illumination angles [36, 39], our strategy provides a somewhat higher frame rate, making it better suited for imaging dynamic samples. Recovering phase information from only 1D gradient images is also possible, where only two complementary illumination angles are required [40, 41], however this tends to be less robust for complex samples because of missing gradient information.

### 2.3. Bead-sample preparation and imaging

Polystyrene beads suspensions (24.7 μm or 45 μm diameter, Polyscience Inc.) were first dried and mixed with a Sylgard 184 base (Dow Corning Corp.) on a microscope slide using a lab spatula. A 10% (w/v) curing agent was then added to the mixture for polymerization. The sample was placed under room temperature for 2 days until complete polymerization. During imaging, a Teflon block was placed under the sample to serve as a bulk scattering medium. Imaging was performed using a 20× /0.42NA objective (Mitutoyo Plan Apo 20×) and 9-plane prism, providing an interplane separation of Δ*z* = 29.5 μm (*n*_sample_ = 1.43, *f*_obj_ = 10 mm).

### 2.4. Chick embryo preparation and imaging

All animal studies were carried out in accordance with the guidelines of the Institutional Animal Care and Use Committee of Boston University. Fertile eggs of While Leghorn (Texas A&M Poultry Science) were incubated at 37.5°C 50% humidity in an egg incubator with automatic turning every 8 hours. Imaging was performed at embryonic day 6 - 11. The top region of the shell and shell membrane were removed to expose the embryo and CAM for imaging. After imaging, the embryos were euthanized by hypothermia by storing the eggs at –15° C. A 6-plane prism and either a 20×/0.42NA objective (Mitutoyo Plan Apo 20×) or a 40× W/0.8NA objective (Olympus UMPLFLN 40× W) were used for imaging, providing an interplane separation of 27.4 μm or 5.55 μm (*n*_sample_ = 1.33, *f*_obj_ = 10 mm or 4.5 mm) with total axial span of 137 μm or 27.8 μm respectively.

## 3. Results

### 3.1. Polystyrene beads

To test our system with more complex samples, we started by imaging mixtures of 24.7 μm polystyrene beads embedded in PDMS. A 20×/0.42NA objective and 9-plane prism provided imaging over a total axial span of 236 μm with an interplane separation of Δ*z* = 29.5 μm. Figure 3(a) shows the raw OBM images from 3 different oblique illumination angles, where both phase-gradient and absorption contrast are entangled. Using Eq. 3 and 4, we can extract pure phase-gradient images along both *x* and *y* axes. The resulting phase-gradient contrast images along the vertical axis across all 9 focal planes are shown in Fig. 3(b). The optical sectioning effect inherent in phase-gradient imaging is apparent, where a single fluorescent bead is mostly apparent in only one or two focal planes. At the same transverse location, different features can be observed at different depths [Fig. 3(c)]. The corresponding phase images at each depth [Fig. 3(d)] are then obtained by integrating the phase-gradient images. Because polystyrene has a higher refractive index than PDMS, the individual beads appear bright against a dark background. We note that some residual background is observable in Fig. 3(d) because the optical sectioning strength of OBM is only moderate, scaling with *z*^−3/2^ with *z* here being defocus distance [32]. In principle this residual background can be removed by deconvolution.

**Fig. 3.**
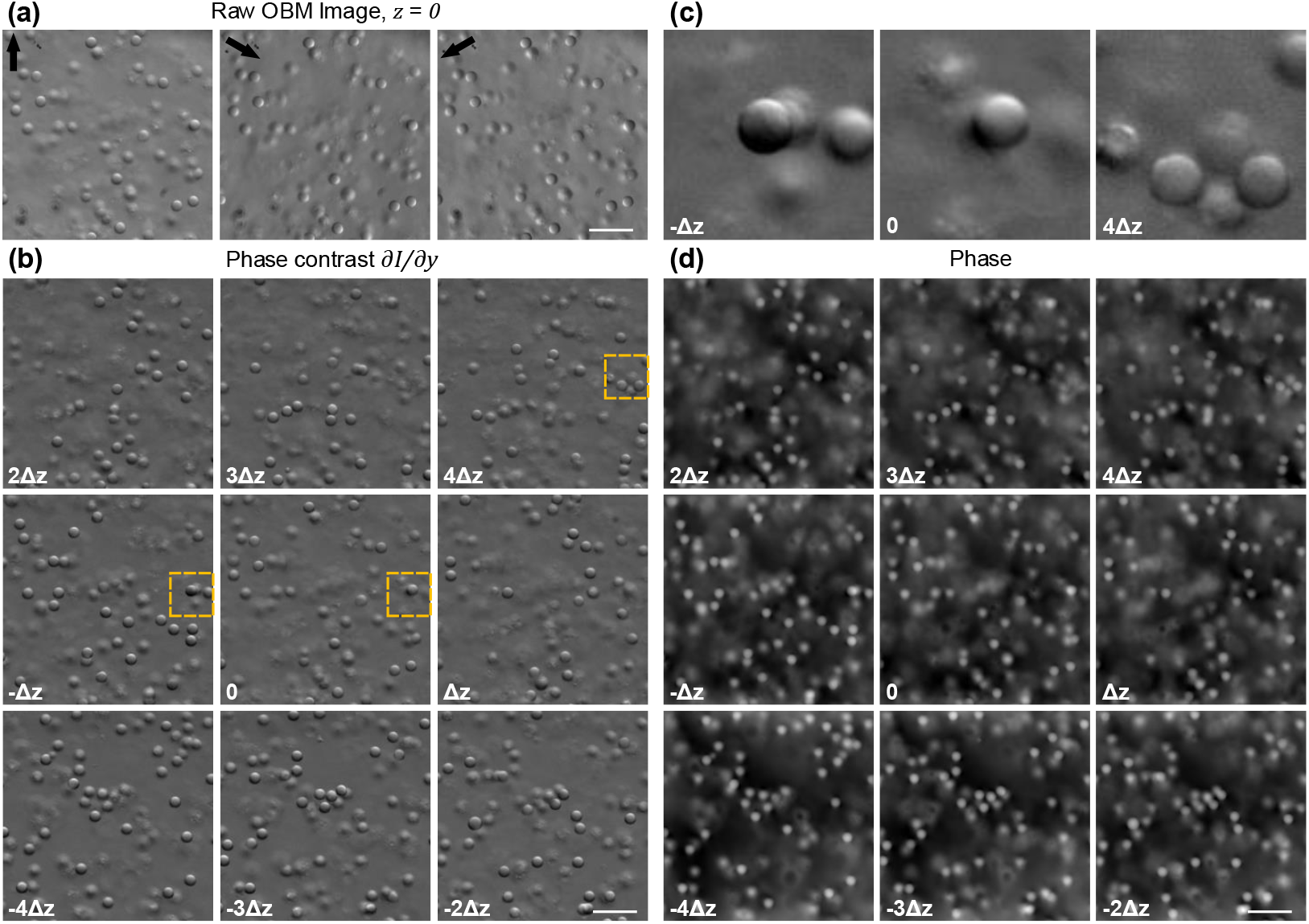
(a) Raw OBM images at depth *z* = 0 obtained from 3 azimuthal back-illumination angles. (b) Synthesized phase-gradient contrast images at 9 different depths. (c) Expanded views over the regions indicated by the yellow boxes in (b) from different depths at Δ*z*, 0, and 4Δ*z*. (d) Reconstructed qualitative phase images from 9 different depths. Δ*z* = 29.5 μm. Scale bar, 100 μm.

### 3.2. Chick Embyro

To highlight the advantage of multifocus OBM for dynamic imaging, we next performed high-speed imaging of chick embryos *in vivo*. An advantage of using a z-splitter prism for multifocus imaging is that different prisms can be readily swapped to prioritize imaging speed or acquisition volume. To prioritize higher imaging speed, we used the 6-plane prism configuration with aspect ratio of 2:3, which allowed us to crop the sensor area-of-interest to 2/3 of the full frame (2048 × 1320 pixels), resulting in 50% faster frame rate. Our acquisition frame rate became then 141 Hz, leading to a net frame rate of 47 Hz for the reconstructed phase images.

With a 20× /0.42NA objective, we first imaged a larger FOV of 546 × 546 × 137 μm^3^ with interplane separation of Δ*z* = 27.4 μm. The resulting phase-gradient images across 6 focal planes are shown in Fig. 4 (a), where a global sample curvature is clearly apparent. With previous implementations of OBM, such curved samples would require additional axial scanning to be fully imaged. Phase images qualitatively corresponding to sample index-of-refraction distributions can be calculated by integration [Fig. 4(b)], also revealing in-focus tissue structures distributed across different focal planes [Fig. 4(b)]. Vascular structures and active blood flow are observable in Visualization 1 and Visualization 2. This volumetric information can be presented more compactly in the form of an extended-depth-of-field (EDOF) image. Various focal stacking techniques are available to do this. Here we used a complex wavelet-based method [42] to generate all-in-focus images from the captured images at different depths. Resulting EDOF phase-gradient and qualitative phase contrast images are shown in Fig. 4(c,d), where the majority of the FOV appears in focus in a single image with relatively high contrast. It should be emphasized that this all-in-focus image does not represent volumetric qualitative or quantitative phase information, but rather simply aids in the visualization of the most prominent features within the imaging volume. A common problem when performing *in vivo* imaging is physiologically induced tissue motion caused by breathing or heart beats. By virtue of the instantaneous nature of our focal-stack acquisition, such motion can be largely accommodated with our mutifocus OBM system. To demonstrate this, we imaged a more mature chick embryo with a readily apparent heart beat. The imaging was performed with a 40 W objective with interplane separation of Δ*z* = 5.5 μm. Figure 5(a,b) shows the 6-plane phase images of CAM at two different time points t = 1.40 s and t = 1.64 s. Tissue displacement is clearly apparent, particularly in the axial direction. Nevertheless, similar structures can be readily followed as they change depths between the time points, with roughly 2Δ*z* axial displacement [Fig. 5(c,d)]. Were it not for multifocus imaging, these structures would be unobservable when out of focus. All-in-focus images from the two time points also exhibit largely similar structures despite axial tissue motion [Fig. 5(e,f)]. Phase-gradient and phase videos of the sample can be found in Visualization 3 and Visualization 4.

**Fig. 4.**
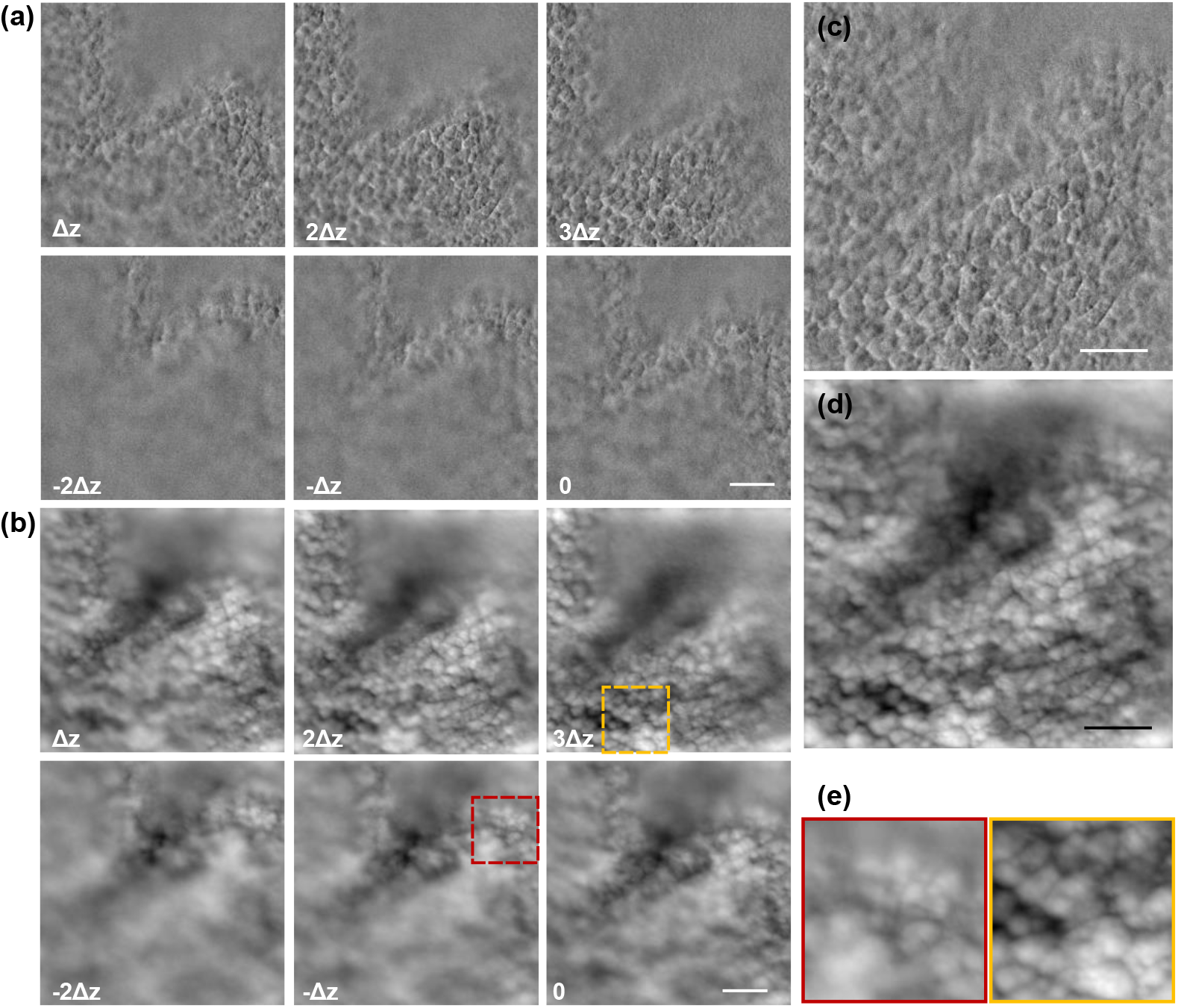
Multifocus OBM applied to *in vivo* imaging of chick CAM. (a) Synthesized phase-gradient contrast images of chick CAM from 6 different depths. (b) Reconstructed qualitative phase images of chick CAM from 6 different depths. (c) A merged all-in-focus phase-gradient contrast image over the imaging volume (−2Δ*z* to 3Δ*z*, total 137 μm axial range). (d) A merged all-in-focus qualitative phase image over the imaging volume. (e) Expanded view from the yellow boxes in (b) at depths Δ*z*, −Δ*z*, and 3Δ*z*. Δ*z* = 27.4 μm. Scale bar, 100 μm.

**Fig. 5.**
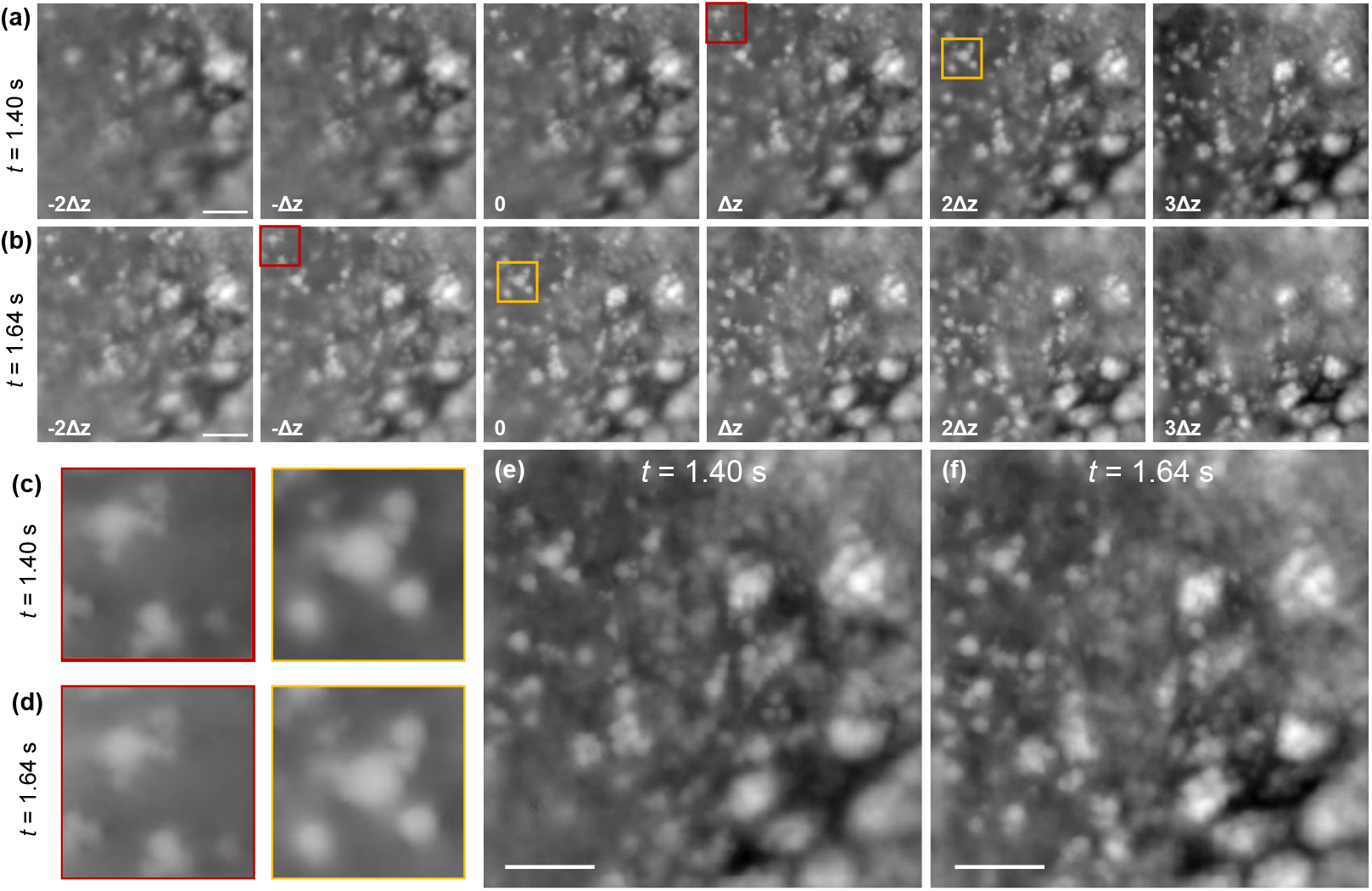
Multifocus OBM is able to accommodate tissue motion during *in vivo* imaging. (a,b) Reconstructed qualitative phase images of chick CAM from 6 different depths at time *t* = 1.40 s and *t* = 1.64 s, respectively. (c,d) Expanded views from the boxed regions in (a) and (b) respectively. (e.f) Merged all-in-focus qualitative phase images from time *t* = 1.40 s and *t* = 1.64 s respectively. Δ*z* = 5.55 μm. All scale bars are 50 μm.

We take advantage of the high frame rate of our multifocus OBM system to *in vivo* image blood flow and track individual red blood cells in 3D. Microvasculature blood-flow imaging is most commonly done with fluorescence angiography [43] or optical coherence tomography [44], where the volumetric imaging speed is generally limited by the fluorescence intensity and/or scanning speed. As an alternative, OBM has allowed *in vivo* label-free imaging of blood flow at the camera native frame rate but only at a single focal plane [35]. Here with a z-splitter prism, 6 focal planes spanning a volume of 241 × 241 × 27.8 μm can be imaged simultaneously, and qualitative multifocus phase reconstruction performed with 3 camera snapshots [Fig. 6(a,b)].

**Fig. 6.**
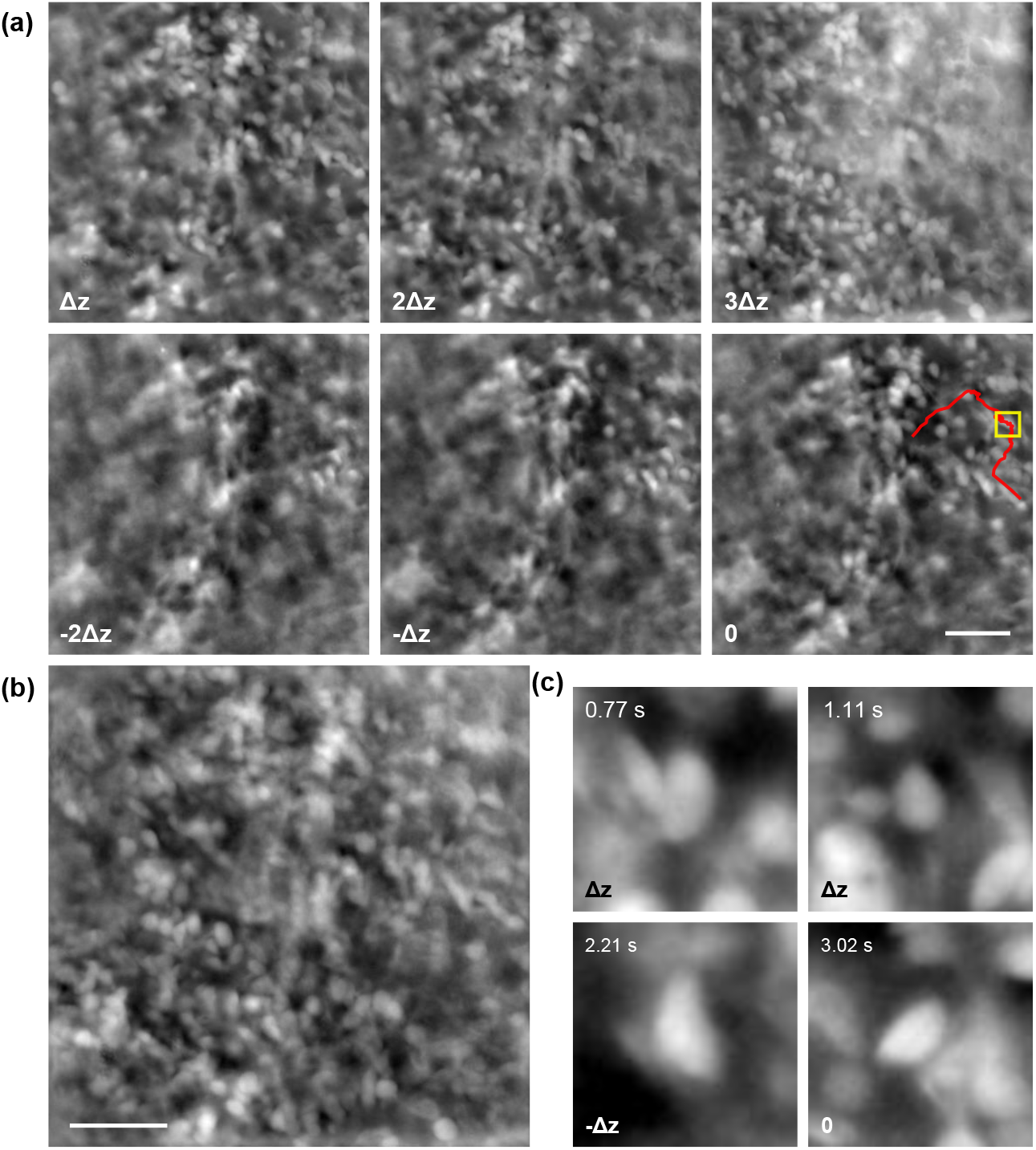
Multifocus OBM applied to chick embryo vasculature imaging and tracking of individual red blood cells in 3D. (a) Reconstructed qualitative phase images of chick embryo vasculature imaged from 6 different depths at time t = 2.26 s. Red trace shows the trajectory of a single red blood cell over a time course of 2.6 s. The position of the cell is identified by the yellow box. (b) A merged all-in-focus image from all images in (a). (c) Images of the same red blood cell tracked in (a) captured at different times from different depths. Δ*z* = 5.55 μm. All scale bars are 50 μm.

The corresponding 6-plane phase-gradient and phase contrast videos are shown in Visualization 5 and Visualization 6]. At a 47 Hz effective frame rate, individual red blood cells can be tracked throughout the imaging volume in 3D. An example trajectory of a cell is marked in red in Fig. 6(a), and in-focus images of this cell at different times are shown in Fig. 6(c) (also see Visualization 7).

## 4. Summary

In summary, we have demonstrated fast multifocus phase imaging by combining OBM [31] with a z-splitter-based microscope [29]. The main advantage of OBM is that it enables phase imaging in arbitrarily thick, scattering samples, which, by definition, are volumetric. Here, with the addition of a z-splitter prism, we are able to capture multiple phase images from within this volume with no penalty in acquisition speed. Because OBM inherently provides optical sectioning [32, 33], it can reveal different axial features within a sample without the need for deconvolution, making it particularly attractive for fast imaging. The resulting combined OBM/z-splitter system offers advantages from both techniques, and yet remains simple, cost-effective, and easy to implement. We were able to demonstrate dynamic phase imaging over a FOV of more than hundreds of micrometers with speeds greater than video-rate, limited here only by our camera frame rate. In principle, this rate could be increased to hundreds or even thousands of hertz by using higher speed cameras that are currently available.

We note that we have confined our interest here to qualitative phase imaging only, with the goal of simply revealing 3D sample structure in a fast and straightforward way. OBM can be extended beyond this to retrieve a full 3D refractive index map of the sample, as has been demonstrated recently by using 3D deconvolution with a simulated point spread function [36]. Our multifocus technique is entirely compatible with this strategy. By obviating the need for axial scanning, our system thus enables the possibility of 3D phase imaging in thick, scattering tissues that is both fast and quantitative.

## Supporting information

Visualization 1

Visualization 2

Visualization 3

Visualization 4

Visualization 5

Visualization 6

Visualization 7

